# Plugging the hunger gap: Organic farming supports more abundant nutritional resources for bees at critical periods

**DOI:** 10.1101/837625

**Authors:** Alexander J. Austin, Lori Lawson-Handley, James D. J. Gilbert

## Abstract

1. Understanding the decline in bee populations and their plant mutualists is of paramount concern for ecosystem health, as well as our future food security. Intensive farming practices are one of the major drivers behind such declines. Organic farming is one of the principal alternatives to conventional practices yet the evidence for its effects are mixed, with some studies showing limited benefits.
2. We conducted bee and floral surveys on 10 paired organic and conventional farms across Yorkshire, UK, to investigate how farming practice influenced the abundance, richness and community composition of bees and flowering plants.
3. Firstly, we found that species richness for flowering plants and bees was similar across organic and conventional farms. Floral composition differed between organic and conventional farms with the greatest differences seen in May and June, whereas bee community composition was similar among farming practices.
4. Secondly, both bee and floral abundance were higher in organic farms. Peaks in floral abundance, and corresponding bee abundance, occurred in particular months, most notably in July, with abundance during the rest of the season being similar across both farming practices.
5. *Synthesis and applications*: Our results suggest that higher floral availability on organic farms corresponds with increased bee abundance. Of particular importance was the higher floral abundance during spring, in the pollinator ‘*hungry gap*’, where floral resources are traditionally scarce. However, conventional farms performed comparably to organic farms across the rest of the season, as well as showing similar levels of species richness, diversity and species composition for both flowering plants and bees. We suggest that targeted management on conventional farms, aimed at boosting floral abundance in the spring, when floral abundance is low, could allow conventional farms to make up the shortfall. Additionally, focusing on increasing the diversity of flowering plants, in terms of both phenology and nutritional composition, for both adult bees and their larvae, could improve bee community diversity across both farming systems.

## Introduction

Insects are in the midst of a global crisis, with unprecedented population reductions being seen across numerous taxa (Sánchez-Bayo & Wyckhuys 2019). Some of the strongest evidence for declines occurs within pollinating insect guilds (Rhodes 2018), and more specifically, from bees, a key pollinator group (Goulson *et al. 2015; Goulson & Nicholls* 2016). Similar patterns are also seen in plants (Niedrist *et al.* 2008), with losses in agricultural habitat particularly prominent (Sotherton & Self 2000). In bees, as with plants, there are multiple drivers involved in their declines (Goulson *et al.* 2015); however, intensive agriculture is widely regarded as one of the largest contributors (Goulson & Nicholls 2016). This loss of bee species, along with other pollinators, is of particular concern as they are essential for the pollination of not only many wild plants (Ollerton *et al.* 2011) but also the majority of commercially important crop species (Aizen *et al.* 2009).

Ecologically-friendly farming methods have the potential to mitigate the effects of intensive agriculture upon pollinator declines and, as such, have received much attention (Kovács-Hostyánszki *et al.* 2017). Agri-environment schemes (AES) are often used to encourage such practices (Batáry *et al.* 2015). Certain AES aim to encourage wildflowers and pollinating insects specifically, including wildflower strips, seed mixes and hedgerow enrichment, many of which have been shown to benefit plant-pollinator communities (Wood *et al.* 2015). Yet AES remain controversial, often benefiting a limited suite of species (Wood *et al.* 2016; MacDonald *et al.* 2019), with doubts on whether they are truly effective for conservation management on farmland (Ansell *et al*. 2016; Riley *et al*. 2017).

Another such route is the adoption of organic farming practices. Organic farming tends to be less intensive, utilising less pesticide and fertiliser (Rigby & Cáceres 2001; Pimentel *et al.* 2005) whilst supporting increased habitat heterogeneity (Fuller *et al.* 2005); attributes which have been shown to benefit bee and plant communities (Rader *et al.* 2014; Basu *et al.* 2016). Many studies have demonstrated benefits in terms of species richness and abundance, for both plants and bees (Gabriel & Tscharntke 2007; Kennedy *et al. 2013)*.

Despite this, studies investigating both organic farming or AES do not show how these patterns shift across a season. Both plant and bee phenology are mutually dependent (Morente-López *et al. 2018)*, an important factor to consider when planning interventions as understanding where troughs in abundance and diversity occur can aid targeted management, allowing farmers to concentrate their efforts during times of shortage. For example, spring is often referred to as the pollinator *‘hungry gap’* on farmland (Nowakowski & Pywell 2016), where floral resources can be scarce. Yet, this is a critical time, with adequate floral resources in this period being essential for emerging bees (Moquet *et al.* 2015) and initial colony growth (Williams *et al.* 2012). However, any differences between these early months and later summer months may be lost when analysed together, or inadequately compensated for with general pollinator management schemes.

Despite the evidence for the positive effects of organic farming, findings are not universal, with results often mixed (Weibull *et al.* 2003; Holzschuh et al. 2008; Rundlöf et al. 2009; Brittain *et al.* 2010). These conflicting findings suggest that other factors may confound effects of farming practice (Roschewitz *et al. 2005)*. Benefits can be seen at the farm scale (Weibull *et al.* 2003), demonstrating the importance of individual farmers in any effects seen within their particular farming system. Farms can also vary greatly within their designation in terms of management practices, intensity, and adoption of AES, which themselves vary in intensity (Wood *et al. 2016)*. These discrepancies between studies highlight the need for more fine-grained studies, conducted across a broader array of farming landscapes than currently exist.

In this study we investigated how organic and conventional farming practices influenced the floral and bee communities, across the season from spring through to late summer. Few studies have specifically looked at how the benefits (or lack thereof) of different farming practices change within the bee flight season (Power *et al.* 2012), despite the importance of phenology in structuring pollinator communities (Encinas-Viso *et al.* 2012) and the obvious effects this may have on the design of certain pollinator management schemes, such as wildflower seed mixes and strips (Nowakowski & Pywell 2016). This temporal aspect is of particular importance for bees, given the importance of the ‘*hungry gap*’ mentioned above (Nowakowski & Pywell 2016). Therefore, investigating the effects of farming practice with a phenological mindset is important for understanding effects of farming at finer temporal scales, and for allowing targeted management interventions that maximise plant-pollinator community benefit. As the effects of organic farming can also be influenced by other factors such as the size of the farm (Fuller *et al.* 2005), the surrounding habitat (Roschewitz *et al.* 2005), and crop type (Tuck *et al.* 2014), we paired conventional and organic farms together to reduce confounding effects as far as possible.

We first predicted that richness and community composition (both floral and bee) would not differ between farming types. In many cases, species richness does not differ between farm types (Power & Stout 2011) and organic farms situated in otherwise intensively farmed landscapes have been shown to have little benefit for bee and plant communities (Brittain *et al.* 2010). Second, as organic farms use more of the crop area for additional floral resources (Pimentel *et al.* 2005), and ‘weed’ species are more common (Romero *et al.* 2008), we predicted that the abundance of flowers and consequently bees would be higher on organic sites when compared to conventional sites. Third, given the ‘*hungry gap*’, we predicted that organic farms would show higher seasonal floral abundance peaks (Morente-López *et al.* 2018), and correspondingly higher seasonal peaks in bee abundance. This study provides (a) valuable information to practitioners about the status of bee-plant communities on differing farm types, (b) insights into where improvements can be made to bolster or improve such communities, and (c) information for a more precise assessment of where key shortages in floral resource occur throughout the bee flight season, allowing for more efficient and targeted interventions for farmland floral enhancement.

## Methods

### Study Sites

Ten matched pairs of conventional and organic farms were recruited in the East Riding of Yorkshire and North Yorkshire, UK. Initial contact was made with farmers via letter, email or telephone and, upon response, face-to-face meetings were arranged to allow both site visits and the exchange of information. Organic farming is relatively uncommon in the East Riding of Yorkshire in comparison to other parts of the UK therefore organic farms were recruited first and then conventional farms local to the organic farms were contacted afterwards. Farms were paired together based on soil type and local environmental conditions (see Fig. S1 and Table S1 for farm pair locations and details) and were paired as closely as possible, in terms of distance. All survey locations within a pair were at least 4 km apart, minimising the chance of any bees caught being able to have foraged at both locations (Knight *et al.* 2005; Zurbuchen *et al.* 2010). Eight of the ten farm pairs were arable or arable/pastoral mixed farms and two pairs were dairy/beef farms. Sample sites on arable/pastoral mixed farms were all located in the arable region of the farm.

### Organism Surveys & Sample Identification

A single field was chosen at each farm within a pair to be the sampling site. Where possible, fields within a pair were matched by crop or, when this was not possible, by crop type. Cereals were the primary crop at all farm sites (barring dairy/beef). Each field within a pair was sampled five times from May to September, 2016, between 09:00 and 16:00 on dry, sunny days with only moderate wind speeds (Forup & Memmott 2005). The order in which each farm within a pair was visited was alternated every sampling session to ensure that, a) farms within pairs were sampled at the same time of day, and b) each farm was sampled during both the morning and afternoon (Power & Stout 2011). An 80 × 1 m transect was walked at a steady pace during each sampling session at each site. The transect consisted of crop, margin and hedgerow habitat types and was ‘E’ shaped with the spine running along the hedgerow/margin and the arms stretching out into the crop. Both spine and arms were 20 × 1 m each. This shape was chosen to reflect the relative land-use of the farm as margins and hedgerows make up comparatively less of the area of a farm than does the crop (Evans *et al.* 2013). By adopting this shape and walking the transect at a steady pace, more time was spent sampling within the crop compared to the hedgerows and margins. The initial location of the transect was chosen using a random number generator where the northern edge of the field was assigned 1, the eastern 2 etc. Subsequent visits then followed a clockwise fashion, to ensure that each start point was random whilst also ensuring each margin was sampled at least once.

Floral surveys were conducted at the beginning of each site visit. Any flowering plants (excluding grasses) along the transect were identified (using Rose & O’Reilly 2006) and the number of floral units taken to provide a measure of floral abundance. A single floral unit was defined as any inflorescence or group of inflorescences that could be navigated without a medium sized bee needing to fly between them (Dicks *et al.* 2002). Flowering plants that were not in flower at the time of sampling were not recorded during that visit.

To sample the bee community, the transect was walked for an hour (total sampling effort) using a stopwatch, catching all bees possible in a sweep net. Upon capture, the stopwatch was paused for sample processing so as to ensure the full hour was spent walking the transect. All captured bees were stored singly in 50 ml falcon tubes, the time of capture recorded (along with any flower they were interacting with at capture), and then placed in a cool bag containing ice packs. Once the transect was complete, all captured samples were placed into a freezer (−20°C) on-site to euthanise and store them. Thawed bees were identified to species level using Falk (2015) and sexed under a light microscope.

### Statistical Analysis

All analyses were performed with R statistical software v3.5 (R Core Team, 2018) and bee and floral data were analysed in the same way. Community indices (species richness, Simpson’s index, and Shannon’s index) were calculated from abundance data per site per sampling visit using the package iNEXT (Hsieh *et al.* 2019). The extrapolated values for all indices were used as species accumulation curves fell short of an asymptote.

Generalised Linear Mixed effect Models (GLMMs) were used to investigate whether farm type influenced bee and floral abundance as well as community indices. Farming type and visit month, along with their interaction, were used as predictors, and farm pair as a random factor in the maximal model. Minimal models were determined by reverse stepwise model selection. GLMMs were also used to investigate whether bee abundance was influenced by the floral abundance on a farm, with bee abundance as the response variable and floral abundance, farm type and visit month, as well as all interactions, as predictor variables.

Community structure was analysed within the package vegan (Oksanen *et al.* 2018) in R. Community similarity of both bees and flowers was investigated using PERMANOVA (ADONIS) of relative species abundances with farm type and visit month, plus their interaction, as predictors. Permutations were restricted to within-farm-pair comparisons only. Nonmetric multidimensional scaling (NMDS) ordination on Bray-Curtis distance matrices was used to visualise differences.

## Results

A total of 1332 bees (780 from organic and 552 from conventional) from 39 species were caught across all farm sites (Table 1), with 4 species unique to conventional sites and 6 to organic sites (Table S2). 67,401 floral units were recorded from 105 flowering plant taxa across farm types (Table 2), with 31 taxa unique to conventional sites and 21 unique to organic sites (Table S3).

**Table 1.**
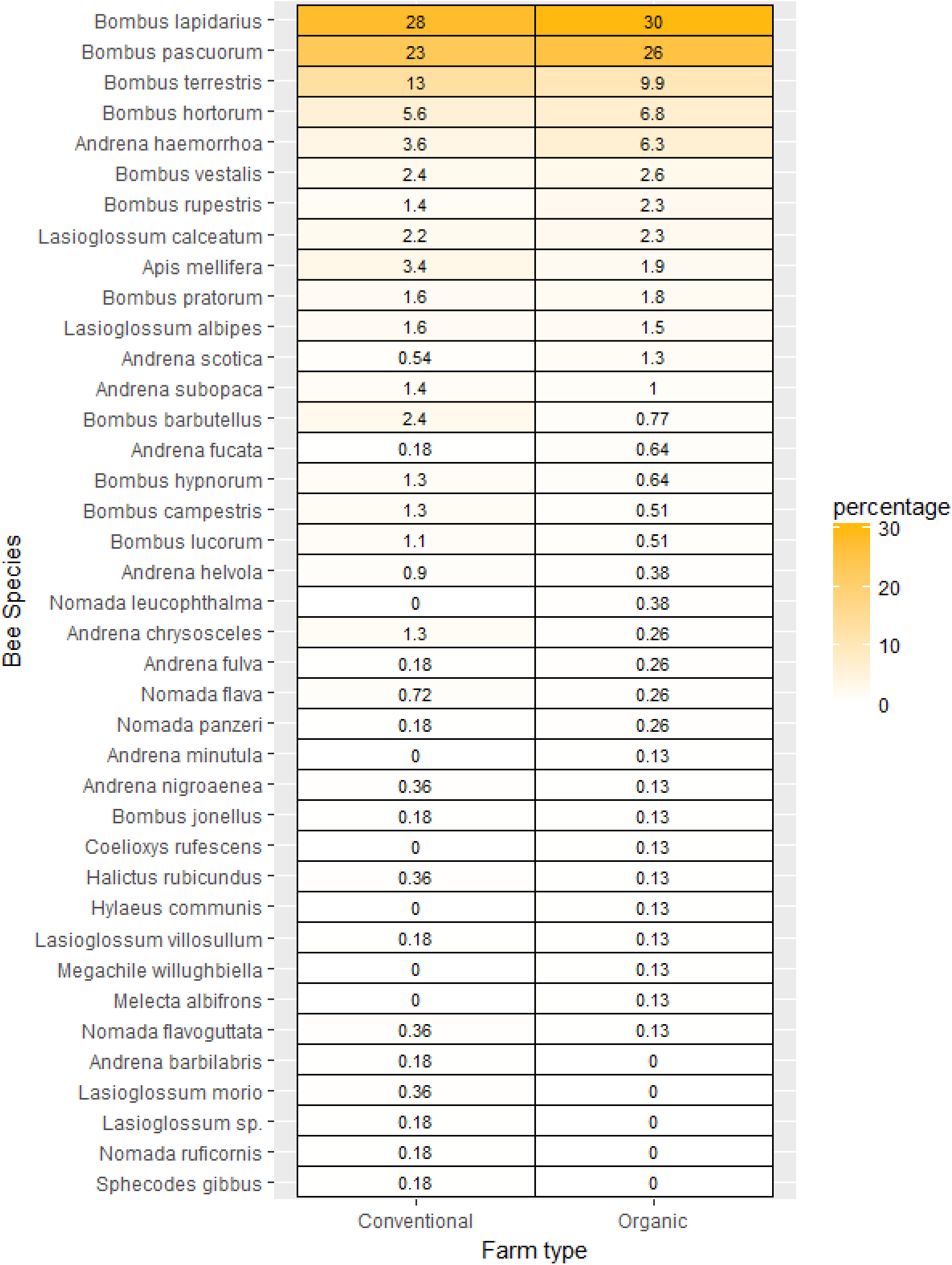
The proportion of each bee species caught on organic and conventional as a percentage of the total number of bees caught per farming type. Ordered by organic farms, highest proportion to lowest.

**Table 2.**
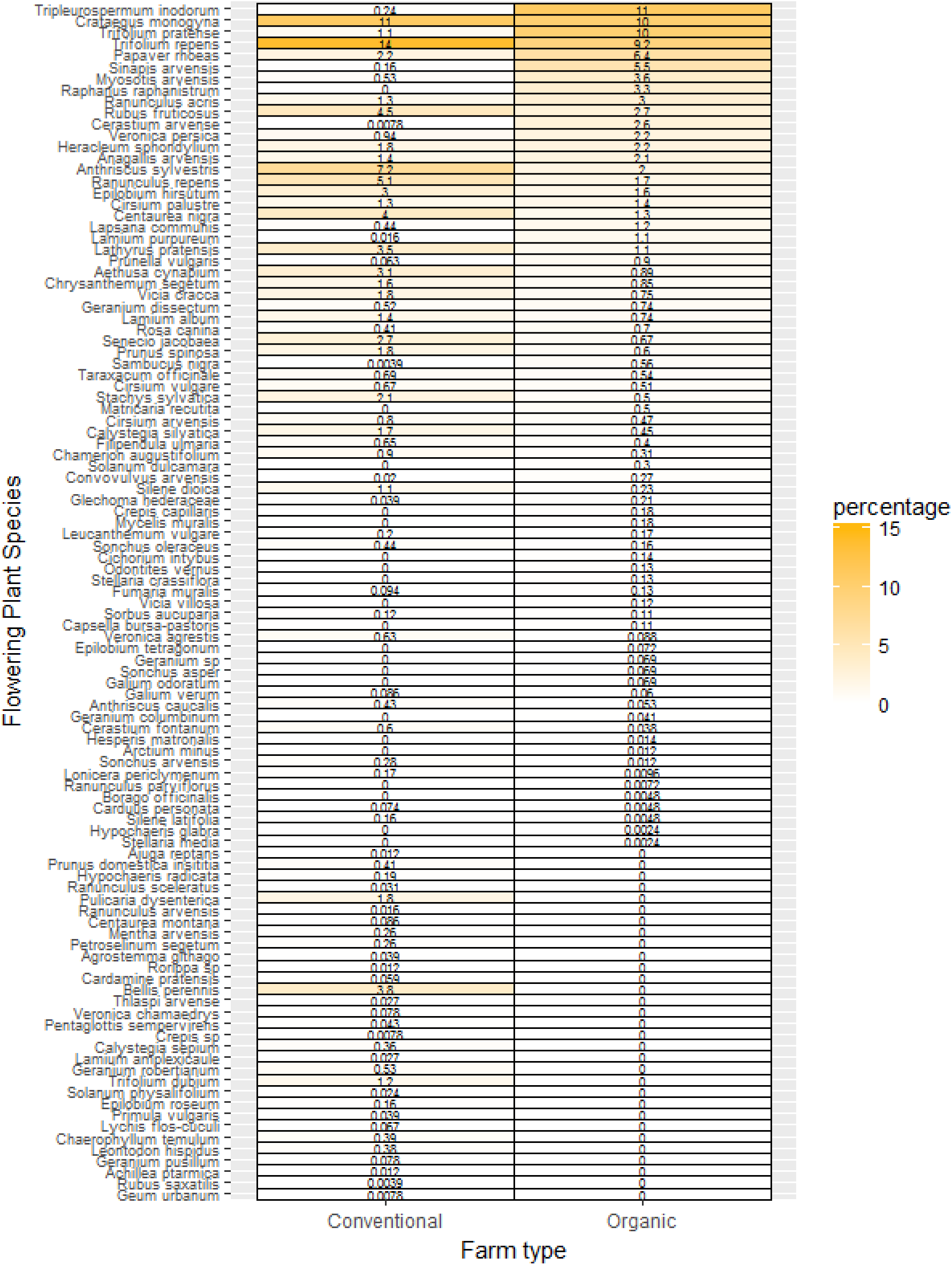
The proportion of each flowering plant taxa recorded on organic and conventional as a percentage of the total number of flowering plants recorded per farming type. Ordered by organic farms, highest proportion to lowest.

### Community diversity indices

The number of bee species caught overall did not differ between farm types (GLMM, dropping effect of farm type. χ^2^_1,7_ = 0.85, p = 0.36). Shifts were seen, however, between months (GLMM, χ^2^_4,6_ = 12.24, p = 0.016), with notable peaks in richness being seen in the months of July and August (Fig. 1a). The same trends were seen in the number of flowering plant species, with farm type having no effect on floral species richness (GLMM, χ^2^_1,7_ = 2.01, p = 0.16) and only the visit month showing a significant effect (χ^2^_4,6_ = 24.50, p<0.001), with higher richness being seen in the summer months (Fig. 1b).

**Fig. 1.**
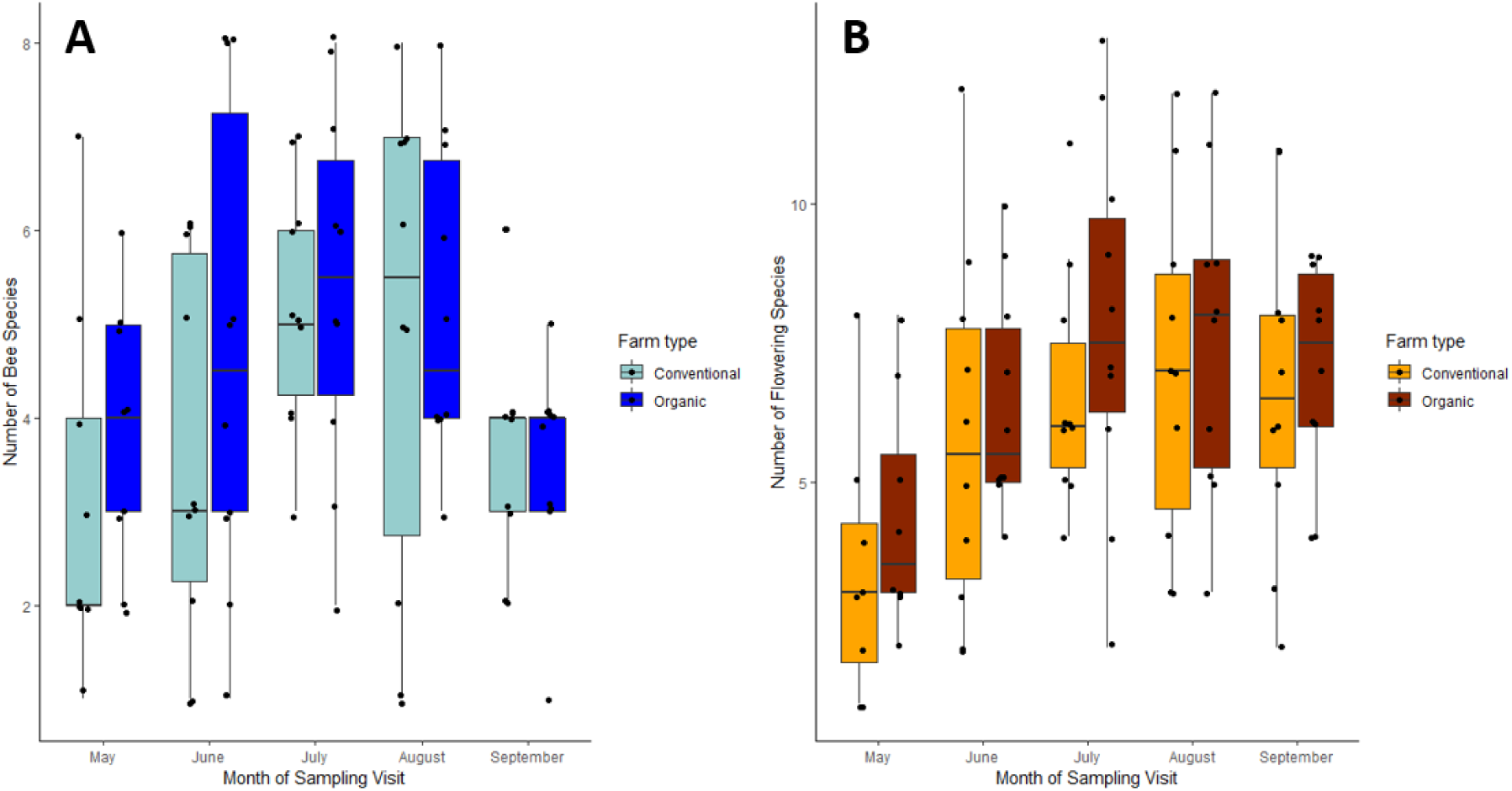
Species richness of bees (A) and flowering plants (B) recorded on organic and conventional farms plotted by sampling month

As the same patterns were seen for both bees and flowering plants for Shannon’s Index and Simpson’s Index, the results for Shannon’s Index only are visualised here. In accordance with the results for richness, Shannon’s Index was not influenced by farm type for either bees (GLMM, χ^2^_1,8_ = 0.88, p = 0.35) or flowering plants (χ^2^_1,8_ = 0.075, p = 0.78), but was significantly influenced by visit month (Bees, χ^2^_1,8_ = 17.55, p = 0.0015, Fig. 2a; Flowering plants, χ^2^_4,7_ = 17.36, p = 0.002, Fig. 2b). For bees, lower index values were seen in September (t_88_ = 3.94, p<0.001) but with flowering plants, May was the lowest diversity month (t_89_ = 9.87, p<0.001), with higher values seen across all other months.

**Fig. 2.**
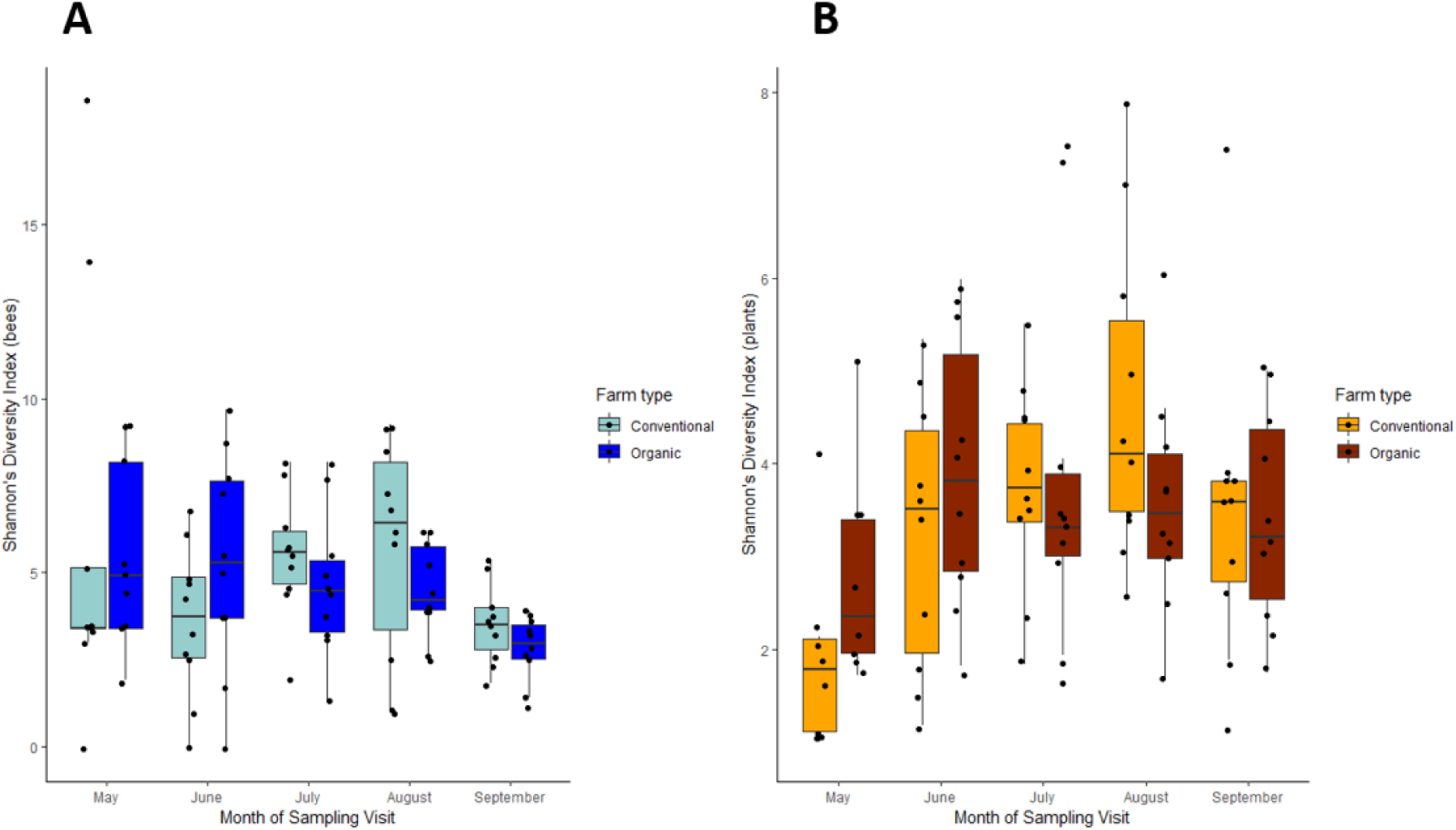
Shannon’s Diversity Index for bees (A) and flowering plants (B) on organic and conventional farms plotted by sample month

The same results were seen for Simpson’s Index for both bees and flowering plants, whereby farm type had no effect (Bees, χ^2^_4,8_ = 2.68, p = 0.1; Flowering plants, χ^2^_1,8_ = 0.06, p = 0.81), with the majority of variation being driven by visit month (Bees, χ^2^_4,8_ = 14.21, p = 0.0067; Flowering plants, χ^2^_4,8_ = 12.77, p = 0.012). Again, as with Shannon’s Index, Simpson’s Index was significantly lower in September for bees (t_91_ = 2.86, p = 0.0042) and significantly lower for flowering plants in May (t_89_ = 9.87, p<0.001).

### Community composition

The bee community composition at farm sites was not influenced by farm type (ADONIS: R^2^ = 0.002, p = 0.986) but was significantly affected by visit month (ADONIS: R^2^ = 0.176, p<0.001), clustering together by the visit month when visualised with NMDS ordination (Fig. 3a,b).

**Fig. 3.**
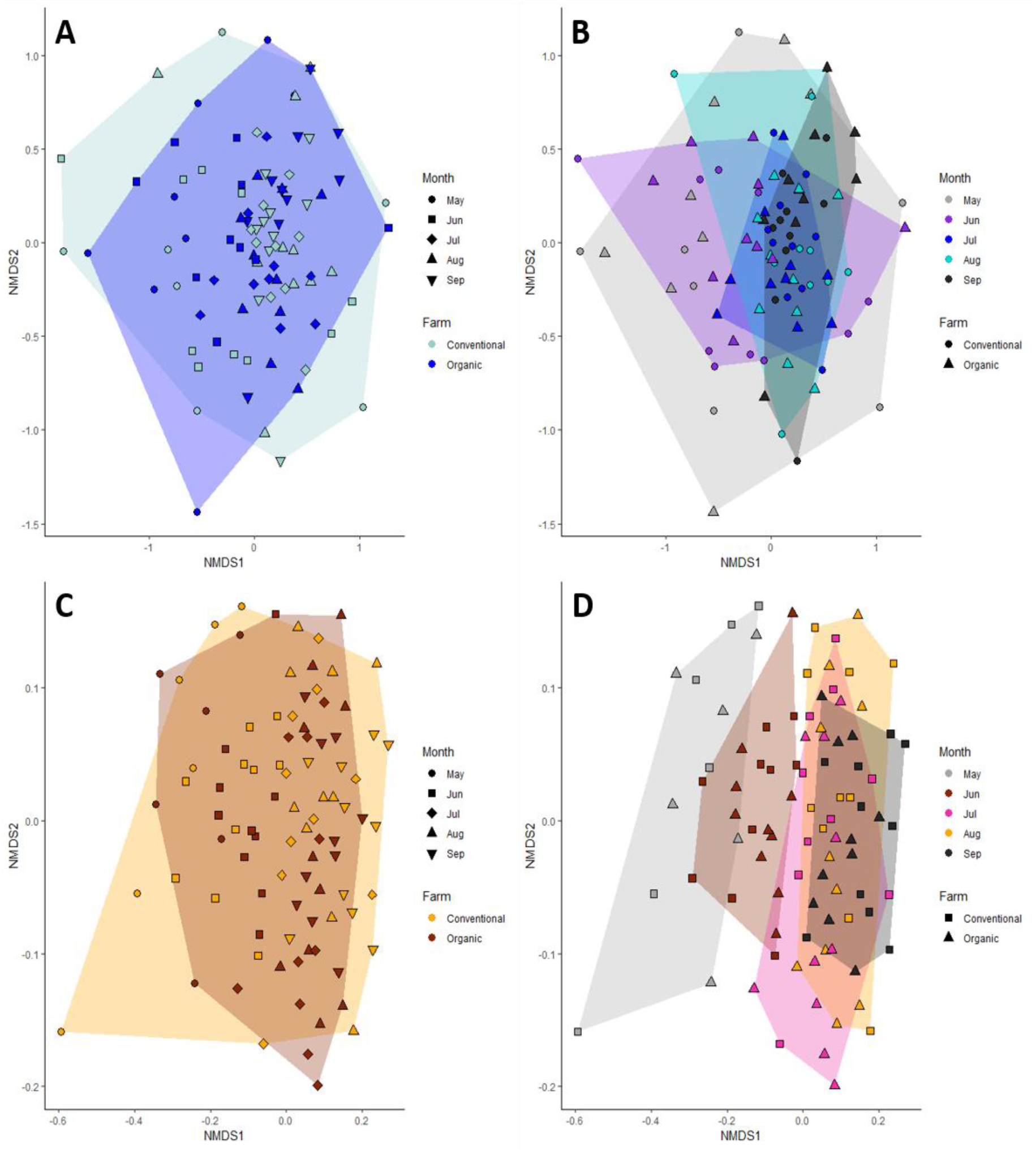
(A, B) NMDS plots both showing the same data: counts of bee species at each site during each sampling month (raw data converted to percentages), but shaded by (A) farm type and sampling month (B). (C, D) NMDS plots both showing the counts of flowering plant species at each site during each sampling month but shaded by farm type (C) and sampling month(D). Note: visit one from one conventional farm was removed as an outlier, its paired organic site was also removed.

The floral community, however, was influenced by both farming type (ADONIS: R^2^ = 0.018, p = 0.0042) and visit month (ADONIS: R^2^ = 0.13, p<0.001). This influence of farming type remained even when one outlying conventional community sample in May was removed (ADONIS: R^2^ = 0.019, p = 0.0044, Fig. S2a). Under NMDS, floral community samples clustered together by visit month, as with the bee communities (Fig. 3c,d). Note that one visit to a conventional site, was removed from ordination analysis (due to only a single specimen being recorded) along with its organic partner. NMDS visualisation with this point included can be seen in Fig. S2. The same conventional site was removed from both bee and floral data and the removal did not influence the outcome of ADONIS analyses.

### Bee & floral abundance

The abundance of bees was influenced by both farming type and visit month with a significant interaction between the two predictors (Generalised Linear Mixed effects Model – GLMM – containing farm type and visit month as predictors with farm pair as a random factor, dropping interaction. χ^2^_4,11_ = 87.727, p<0.001). Overall, more bees were caught on organic farms than on conventional farms (Fig. 4a) with this difference primarily being driven by the months of July and August (Fig. 4b).

**Fig. 4.**
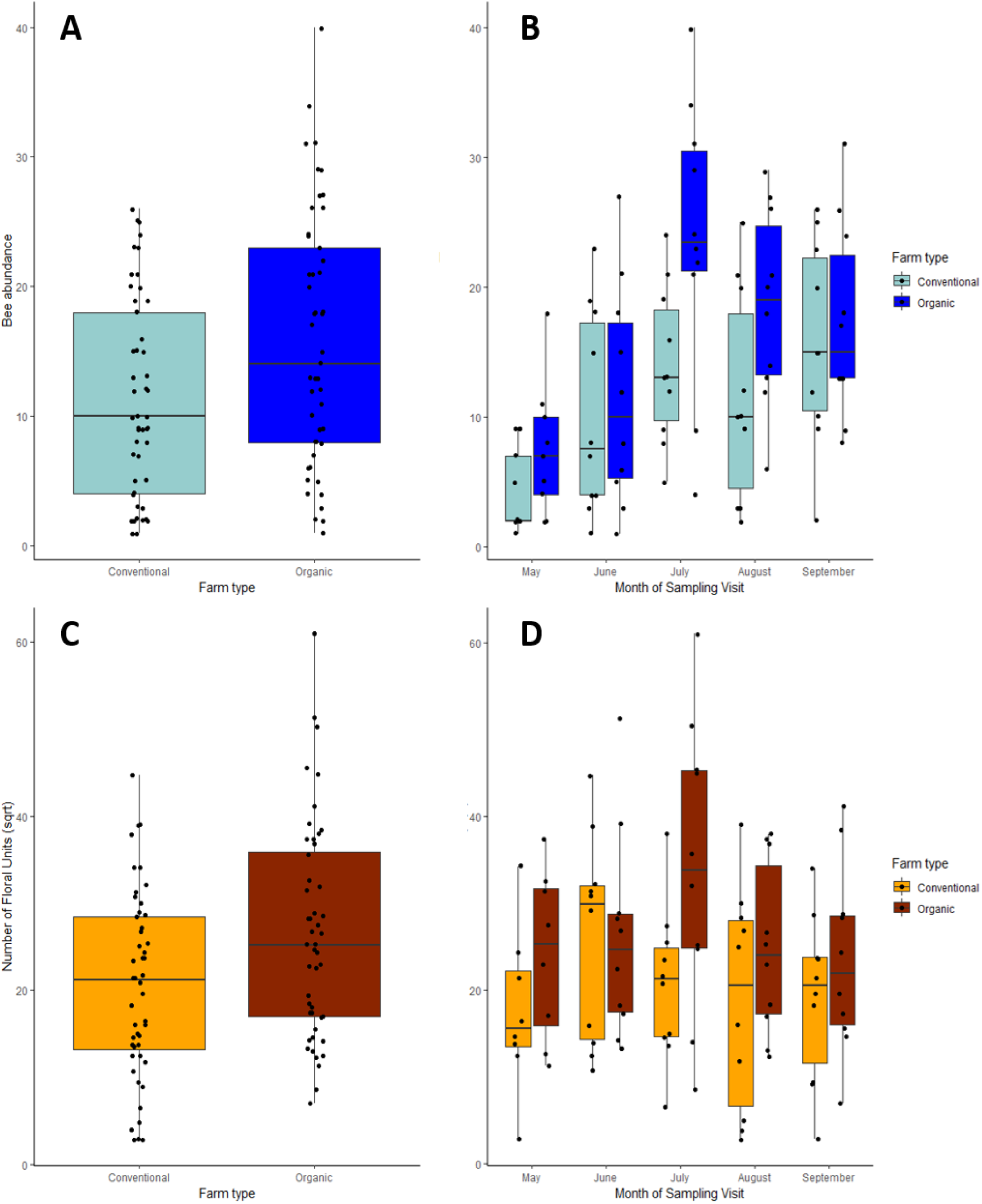
Abundance of bees and flowers recorded on farm sites. Number of bees recorded on organic and conventional farms in totality (A), and within each visit month separately (B). Number of floral units (square root transformed) recorded on organic and conventional farms in totality (C), and within each visit month separately (D).

There was a significant interaction between farm type and visit month when analysing the floral abundance (GLMM, dropping the interaction. χ^2^_7,11_ = 2312.8, p<0.001). Overall, there was a higher number of floral units (greater floral abundance) recorded on organic farms (Fig. 4c) and, as with bee abundance, floral abundance was higher on organic farms in the summer months of July and August, as well as in May (Fig. 4d).

There was a significant 3-way interaction (all predictors) when modelling bee abundance against floral abundance (GLMM, χ^2^_4,21_ = 10.71, p = 0.03), with bee abundance being positively associated with floral abundance to different degrees based on month and farm type (Fig. 5a-e,f). However, it seems clear that the month of May is driving this interaction (Fig. 5a), with bee abundance being negatively associated with floral abundance on organic sites but not on conventional sites.

**Fig. 5.**
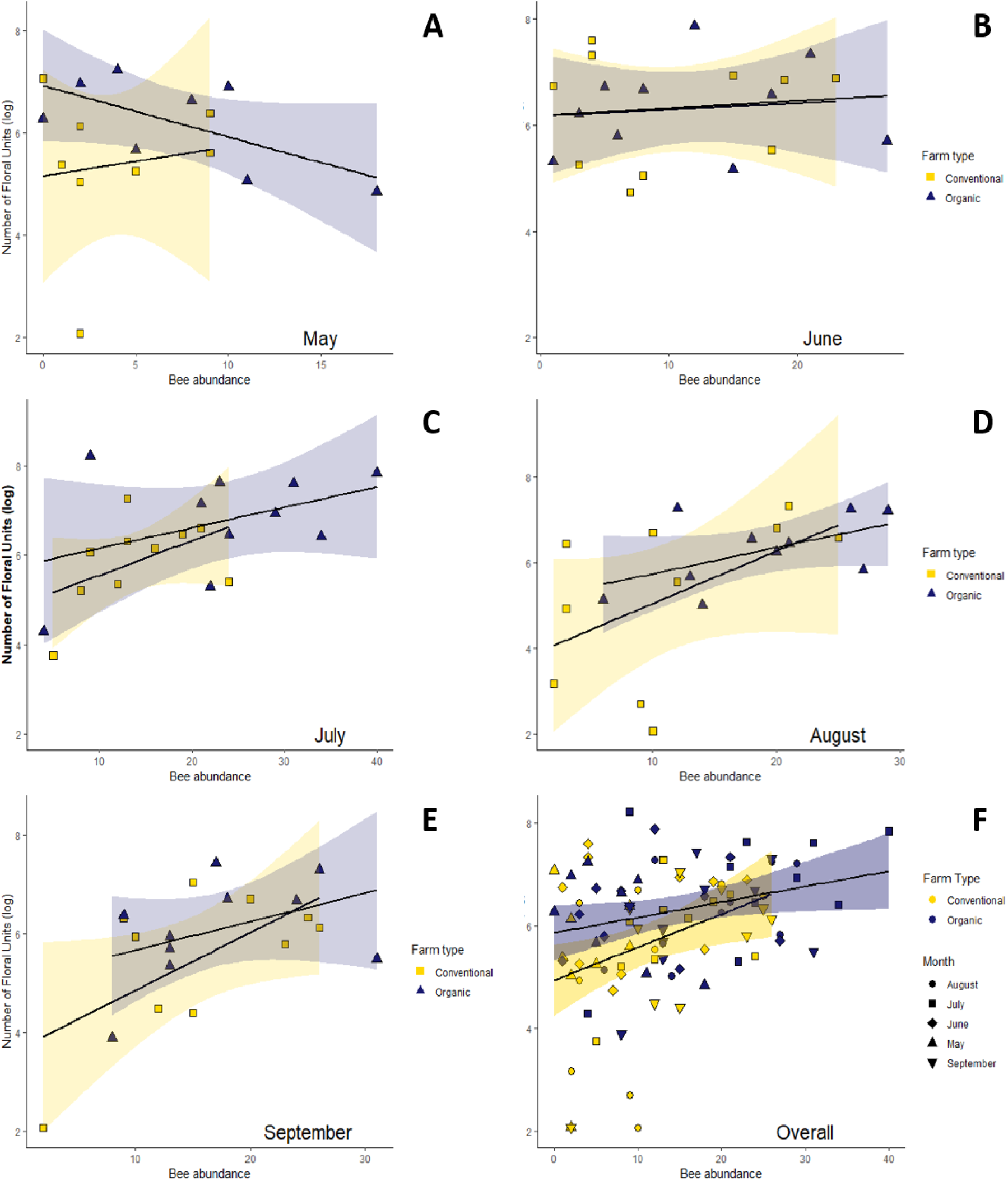
(A-E) the number of bees and the corresponding number of floral units recorded on conventional and organic farms plotted by sampling month. Each point represents a single farm. (F) The total number of bees and corresponding floral units recorded on organic and conventional sites pooled across sampling months. Each point represents a single month at a single farm.

## Discussion

As predicted, organic farms supported higher seasonal and overall floral and bee abundances than conventional farms, though these increases were driven by particular months (e.g. May, July; Fig. 4). Equally, we found no difference in diversity or richness of flowering plants or bees between farming practices (Fig. 1, 2), despite floral community composition differing between farm types (Fig. 3). Overall, we found that phenology and not farming practice had the strongest influence on bee and flowering plant communities in terms of variation in abundances, diversity and community structure. Whilst the benefits of organic farming have received much attention (Hole *et al.* 2005) the relative similarities between farming practices found here suggests that with focussed management interventions, differences in abundance between farm types could be reduced.

Perhaps unsurprisingly, organic farms supported greater abundances of bees and flowering plants, as has been shown previously (Bengtsson *et al.* 2005; Rundlöf et al. 2008, 2009). Interestingly however, the higher abundances in organic farms only occurred in certain months. Organic farms showed the highest abundance relative to conventional farms in July, in both bees and flowering plants, with modest differences in May and August; however, during June and September conventional farms were comparable. May is of particular interest, as spring is often a pollinator ‘*hungry gap*’ on farmland (Nowakowski & Pywell 2016) traditionally linked with poor floral availability. This period is critically important for bees and our results suggest that organic sites may be providing better support during this crucial time. This also suggests that spring should be a focus period for improvement on conventional farms, whereby farmers focus on ensuring spring flowering species in management interventions such as seed mixes, wildflowers strips and hedgerow enrichment.

Increases in bee numbers were related to higher floral unit abundance overall (Fig. 3). As organic farms tend to support high floral abundances (Batáry *et al.* 2013) and, broadly speaking, more flowers tends to mean more bees (Biesmeijer *et al.* 2006; Potts et al. 2010) it could be said that organic farms support more bees. However, this positive association does not simply apply to organic farms; the same is true for conventional sites (Fig. 3; Power & Stout 2011), suggesting conventional farms will also be able to support more bees, if floral availability is improved. Moreover, the increase in bee abundance on organic farms could also be driven by additional factors such as more nesting habitat or reduced exposure to harmful pesticides (Goulson *et al. 2015)*. Interestingly, bee abundance on organic farms in May was negatively related to flower abundance (Fig. 3a), unlike previously identified trends (Gabriel *et al.* 2013; Kennedy *et al.* 2013). This pattern may reflect non-floral resources: without appropriate nesting sites or overwintering sites bee populations can also struggle (Kells & Goulson 2003). Alternatively, with floral resources scarce during the hungry gap, individual bees may spend more time foraging and are therefore encountered more frequently when sampling. In summer months, when availability is high, more bees can be supported and therefore more bees are caught during sampling.

The bee communities found on organic and conventional farms were similar, running counter to some studies (Holzschuh *et al.* 2007, 2010) but agreeing with others (Brittain *et al.* 2010). It is possible that the positive benefits of organic farming may only be apparent when such land makes up a threshold proportion of the agricultural landscape (Brittain *et al.* 2010). As organic farms in Yorkshire are relatively isolated, and far fewer in number than in other parts of the UK, this could explain the uniformity in bee community structure in this case. Equally, as the majority of conventional farms in this study were involved in AES (either ELS or HLS) differences could be expected to be smaller in comparison to their more intensive counterparts (Albrecht et al. 2007)

Unlike with bees, we found that farming practice did influence floral communities, whereby (a) a significant difference in the flowering plant community was seen between organic and conventional sites, and (b) organic sites were more similar to each other than their conventional counterparts (Figs. 3, S2). Although they do share floral species, organic and conventional farms have been shown to support different plant taxa also (Hawes *et al.* 2010). Organic farms support more non-crop species (Gabriel & Tscharntke 2007), which may go some way to explaining the community differences. The bias towards unique flowering plants on organic farms was not found in this study (Table S3), yet the fact that different farming practices supported differing flowering plant species yet similar bee species is interesting. The vast majority of the bees caught in this study were polylectic, i.e. plant generalists, which tend to dominate agricultural landscapes (Ahrenfeldt *et al.* 2019). This generality may help to explain the similarity in bee communities despite the floral differences. Despite the difference seen in the floral communities between farm types it is important to note that the differences seen between months were far greater, suggesting that differences in farmland floral communities during the bee season are affected more by phenological drivers than by farming practice *per se*, as stated above. Therefore, understanding the effects of phenology should not be overlooked in farmland bee-plant communities as it likely has a stronger influence on the health of such systems than the farming practice itself. As such, a focus on ensuring floral availability across the flight season may be just as important to bee communities as a focus on maintaining floral diversity by altering farming practices *per se*.

Despite the differences seen in flowering plant community structure across sampling months, there was no difference in *overall* species richness or diversity between farming practices. Previous studies show mixed results for both plants and bees, with some showing higher levels of richness and diversity on organic farms (Gabriel & Tscharntke 2007; Kennedy *et al.* 2013) but others not (Gibson *et al.* 2007; Brittain et al. 2010). The effects of farming practice can be heavily influenced by the composition of the surrounding landscape (Roschewitz *et al.* 2005; Brittain et al. 2010). Although statistically there was no difference between farming practices, it is worth noting that floral richness and diversity do seem to fluctuate across months, with May in particular showing lower diversity and species richness, coinciding with that of bees (Fig. 1, 2), highlighting the ‘*hungry gap*’ on farmland.

We found little difference in the richness and diversity of flowering plants, or bees, between farming practices (Fig. 4b, 5b), however, what is clear is that spikes in floral abundance correspond with spikes in bee abundance (Fig. 4b, d). These gluts in floral abundance may be useful sources of nutrition but bee communities require floral constancy across the season in order to support them throughout their life cycles (Goulson & Nicholls 2016); something both farming systems could improve upon.

Organic farming appears to support greater abundances of floral resources and bee pollinators, yet, in line with some previous studies, does not seem to support higher levels of diversity in either group (Gibson *et al.* 2007; Brittain et al. 2010). The higher floral abundances coincide with higher abundance in bees, and these levels are substantially higher during the months of May and July on organic farms. However, floral and bee communities on both farm types are similar and conventional sites may be able to plug the gaps by creating flower-rich habitats (Carvell *et al.* 2007). Additionally, as organic farms only showed increases in floral resources during specific months, targeted management towards these periods of scarcity (Nowakowski & Pywell 2016) on conventional farms may allow then to catch up. However, as we have seen from our study, along with others (Power & Stout 2011), increased floral and bee abundance does not correspond to an increase in diversity, therefore “more flowers equals more bees” could be seen as a rather simplistic view of a complex issue. As such, ensuring that management practices for both farming methods take into account diversity and quality (in terms of nutrition) of floral resources will likely support a greater number and diversity of not only flowers (Gibson *et al.* 2007) but bees also (Filipiak *et al.* 2017). Additionally, individual farmers can make a difference, as many of these effects can be seen at the farm scale (Weibull *et al.* 2003), especially for solitary bee species, which respond to relatively small management improvements at smaller scales (Steffan-Dewenter *et al.* 2002). Our findings reinforce the idea that biodiversity support in agriculture is more complex than simply organic or conventional, with other parameters also playing a role (Kennedy *et al. 2013)*. Perhaps then, such pigeon-holing may be of little benefit, and encouraging farmers to adopt wildlife-friendly farming practices (Pywell *et al.* 2015) irrespective of their designation should be the focus, as well as communicating the benefits of such methods (Gabriel *et al.* 2013; Pywell *et al.* 2015) directly to the end user i.e. the farmers themselves. Both making farmers aware of such benefits, as well as continuing to research new ways to improve management schemes, should be a priority. For example, pollinator seed mixes have been shown to benefit certain species (Blackmore & Goulson 2014), yet provide little support during certain periods (Pywell *et al.* 2011) or for some groups, such as solitary bees (Wood *et al. 2016)*. Additionally, an increased focus on the nutritional *quality* of such resources should be used to inform the inclusion of particular flowers into such seed mixes. The importance of quality nutrition to bee health is becoming more and more apparent (Fowler *et al.* 2016; Nicholls & Hempel de Ibarra 2016), with pollen in particular being a highly variable (Roulston & Cane 2000) yet critical resource for bee development (Donkersley *et al.* 2017; Filipiak 2019). Improvements to these schemes, coupled with targeted management, could not only allow for conventional farms to plug the gap in floral and bee abundances seen in comparison to organic farms, but also lead to improvements in the diversity of bee (and wider pollinator) communities across all farm practices.

## References

Ahrenfeldt, E.J., Kollmann, J., Madsen, H.B., Skov-Petersen, H. & Sigsgaard, L. (2019). Generalist solitary ground-nesting bees dominate diversity survey in intensively managed agricultural land. Journal of Melittology, 1–12.

Aizen, M.A., Garibaldi, L.A., Cunningham, S.A. & Klein, A.M. (2009). How much does agriculture depend on pollinators? Lessons from long-term trends in crop production. Ann. Bot., 103, 1579–1588.

Ansell, D., Freudenberger, D., Munro, N. & Gibbons, P. (2016). The cost-effectiveness of agri-environment schemes for biodiversity conservation: A quantitative review. Agric. Ecosyst. Environ., 225, 184–191.

Basu, P., Parui, A.K., Chatterjee, S., Dutta, A., Chakraborty, P., Roberts, S., et al. (2016). Scale dependent drivers of wild bee diversity in tropical heterogeneous agricultural landscapes. Ecol. Evol.

Batáry, P., Dicks, L.V., Kleijn, D. & Sutherland, W.J. (2015). The role of agri-environment schemes in conservation and environmental management. Conserv. Biol., 29, 1006–1016.

Batáry, P., Sutcliffe, L., Dormann, C.F. & Tscharntke, T. (2013). Organic farming favours insect-pollinated over non-insect pollinated forbs in meadows and wheat fields. PLoS One, 8, e54818.

Bengtsson, J., Ahnström, J. & Weibull, A.-C. (2005). The effects of organic agriculture on biodiversity and abundance: a meta-analysis. J. Appl. Ecol., 42, 261–269.

Biesmeijer, J.C., Roberts, S.P.M., Reemer, M., Ohlemüller, R., Edwards, M., Peeters, T., et al. (2006). Parallel declines in pollinators and insect-pollinated plants in Britain and the Netherlands. Science, 313, 351–354.

Blackmore, L.M. & Goulson, D. (2014). Evaluating the effectiveness of wildflower seed mixes for boosting floral diversity and bumblebee and hoverfly abundance in urban areas. Insect Conserv. Divers., 7, 480–484.

Brittain, C., Bommarco, R., Vighi, M., Settele, J. & Potts, S.G. (2010). Organic farming in isolated landscapes does not benefit flower-visiting insects and pollination. Biol. Conserv., 143, 1860–1867.

Carvell, C., Meek, W.R., Pywell, R.F., Goulson, D. & Nowakowski, M. (2007). Comparing the efficacy of agri-environment schemes to enhance bumble bee abundance and diversity on arable field margins. J. Appl. Ecol., 44, 29–40.

Dicks, L.V., Corbet, S.A. & Pywell, R.F. (2002). Compartmentalization in plant–insect flower visitor webs. J. Anim. Ecol., 71, 32–43.

Donkersley, P., Rhodes, G., Pickup, R.W., Jones, K.C., Power, E.F., Wright, G.A., et al. (2017). Nutritional composition of honey bee food stores vary with floral composition. Oecologia.

Encinas-Viso, F., Revilla, T.A. & Etienne, R.S. (2012). Phenology drives mutualistic network structure and diversity. Ecol. Lett., 15, 198–208.

Evans, D.M., Pocock, M.J.O. & Memmott, J. (2013). The robustness of a network of ecological networks to habitat loss. Ecol. Lett., 16, 844–852.

Falk, S.J. (2015). Field guide to the bees of Great Britain and Ireland. British Wildlife Publishing.

Filipiak, M. (2019). Key pollen host plants provide balanced diets for wild bee larvae: A lesson for planting flower strips and hedgerows. J. Appl. Ecol.

Filipiak, M., Kuszewska, K., Asselman, M., Denisow, B., Stawiarz, E., Woyciechowski, M., et al. (2017). Ecological stoichiometry of the honeybee: Pollen diversity and adequate species composition are needed to mitigate limitations imposed on the growth and development of bees by pollen quality. PLoS One, 12, e0183236.

Forup, M.L. & Memmott, J. (2005). The Restoration of Plant–Pollinator Interactions in Hay Meadows. Restor. Ecol., 13, 265–274.

Fowler, R.E., Rotheray, E.L. & Goulson, D. (2016). Floral abundance and resource quality influence pollinator choice. Insect Conserv. Divers.

Fuller, R.J., Norton, L.R., Feber, R.E., Johnson, P.J., Chamberlain, D.E., Joys, A.C., et al. (2005). Benefits of organic farming to biodiversity vary among taxa. Biol. Lett., 1, 431–434.

Gabriel, D., Sait, S.M., Kunin, W.E. & Benton, T.G. (2013). Food production vs. biodiversity: comparing organic and conventional agriculture. J. Appl. Ecol., 50, 355–364.

Gabriel, D. & Tscharntke, T. (2007). Insect pollinated plants benefit from organic farming. Agric. Ecosyst. Environ., 118, 43–48.

Gibson, R.H., Pearce, S., Morris, R.J., Symondson, W. & Memmott, J. (2007). Plant diversity and land use under organic and conventional agriculture: a whole-farm approach. J. Appl. Ecol., 44, 792–803.

Goulson, D. & Nicholls, E. (2016). The canary in the coalmine; bee declines as an indicator of environmental health. Sci. Prog., 99, 312–326.

Goulson, D., Nicholls, E., Botías, C. & Rotheray, E.L. (2015). Bee declines driven by combined stress from parasites, pesticides, and lack of flowers. Science, 347, 1255957.

Hawes, C., Squire, G.R., Hallett, P.D., Watson, C.A. & Young, M. (2010). Arable plant communities as indicators of farming practice. Agric. Ecosyst. Environ., 138, 17–26.

Hole, D.G., Perkins, A.J., Wilson, J.D., Alexander, I.H., Grice, P.V. & Evans, A.D. (2005). Does organic farming benefit biodiversity? Biol. Conserv., 122, 113–130.

Holzschuh, A., Steffan-Dewenter, I., Kleijn, D. & Tscharntke, T. (2007). Diversity of flower-visiting bees in cereal fields: effects of farming system, landscape composition and regional context. J. Appl. Ecol., 44, 41–49.

Holzschuh, A., Steffan-Dewenter, I. & Tscharntke, T. (2008). Agricultural landscapes with organic crops support higher pollinator diversity. Oikos, 117, 354–361.

Holzschuh, A., Steffan-Dewenter, I. & Tscharntke, T. (2010). How do landscape composition and configuration, organic farming and fallow strips affect the diversity of bees, wasps and their parasitoids?. J. Anim. Ecol., 79, 491–500.

Hsieh, T.C., Ma, K.H. & Chao, A. (2019). iNEXT: iNterpolation and EXTrapolation for species diversity. R package version 2.0. 19.

Kells, A.R. & Goulson, D. (2003). Preferred nesting sites of bumblebee queens (Hymenoptera: Apidae) in agroecosystems in the UK. Biol. Conserv., 109, 165–174.

Kennedy, C.M., Lonsdorf, E., Neel, M.C., Williams, N.M., Ricketts, T.H., Winfree, R., et al. (2013). A global quantitative synthesis of local and landscape effects on wild bee pollinators in agroecosystems. Ecol. Lett., 16, 584–599.

Knight, M.E., Martin, A.P., Bishop, S., Osborne, J.L., Hale, R.J., Sanderson, R.A., et al. (2005). An interspecific comparison of foraging range and nest density of four bumblebee (Bombus) species. Mol. Ecol., 14, 1811–1820.

Kovács-Hostyánszki, A., Espíndola, A., Vanbergen, A.J., Settele, J., Kremen, C. & Dicks, L.V. (2017). Ecological intensification to mitigate impacts of conventional intensive land use on pollinators and pollination. Ecol. Lett.

MacDonald, M.A., Angell, R., Dines, T.D., Dodd, S., Haysom, K.A., Hobson, R., et al. (2019). Have Welsh agri-environment schemes delivered for focal species? Results from a comprehensive monitoring programme. J. Appl. Ecol., 49, 871.

Moquet, L., Mayer, C., Michez, D., Wathelet, B. & Jacquemart, A.-L. (2015). Early spring floral foraging resources for pollinators in wet heathlands in Belgium. J. Insect Conserv., 19, 837–848.

Morente-López, J., Lara-Romero, C., Ornosa, C. & Iriondo, J.M. (2018). Phenology drives species interactions and modularity in a plant – flower visitor network. Sci. Rep., 8, 9386.

Nicholls, E. & Hempel de Ibarra, N. (2016). Assessment of pollen rewards by foraging bees. Funct. Ecol.

Niedrist, G., Tasser, E., Lüth, C., Dalla Via, J. & Tappeiner, U. (2008). Plant diversity declines with recent land use changes in European Alps. Plant Ecol., 202, 195.

Nowakowski, M. & Pywell, R. (2016). Habitat Creation and Management for Pollinators. Centre for Ecology & Hydrology, Wallingford, UK.

Oksanen, J., Blanchet, F.G., Friendly, M., Kindt, R., Legendre, P., McGlinn, D., et al. (2018). vegan: Community Ecology Package. R package version 2.5-2. 2018.

Ollerton, J., Winfree, R. & Tarrant, S. (2011). How many flowering plants are pollinated by animals? Oikos.

Pimentel, D., Hepperly, P., Hanson, J., Douds, D. & Seidel, R. (2005). Environmental, Energetic, and Economic Comparisons of Organic and Conventional Farming Systems. Bioscience, 55, 573–582.

Potts, S.G., Biesmeijer, J.C., Kremen, C., Neumann, P., Schweiger, O. & Kunin, W.E. (2010). Global pollinator declines: trends, impacts and drivers. Trends Ecol. Evol., 25, 345–353.

Power, E.F., Kelly, D.L. & Stout, J.C. (2012). Organic farming and landscape structure: effects on insect-pollinated plant diversity in intensively managed grasslands. PLoS One, 7, e38073.

Power, E.F. & Stout, J.C. (2011). Organic dairy farming: impacts on insect–flower interaction networks and pollination. J. Appl. Ecol., 48, 561–569.

Pywell, R.F., Heard, M.S., Woodcock, B.A., Hinsley, S., Ridding, L., Nowakowski, M., et al. (2015). Wildlife-friendly farming increases crop yield: evidence for ecological intensification. Proceedings of the Royal Society B: Biological Sciences, 282, 20151740.

Pywell, R.F., Meek, W.R., Hulmes, L., Hulmes, S., James, K.L., Nowakowski, M., et al. (2011). Management to enhance pollen and nectar resources for bumblebees and butterflies within intensively farmed landscapes. J. Insect Conserv., 15, 853–864.

Rader, R., Birkhofer, K., Schmucki, R., Smith, H.G., Stjernman, M. & Lindborg, R. (2014). Organic farming and heterogeneous landscapes positively affect different measures of plant diversity. J. Appl. Ecol., 51, 1544–1553.

Rhodes, C.J. (2018). Pollinator decline – an ecological calamity in the making? Sci. Prog., 101, 121–160.

Rigby, D. & Cáceres, D. (2001). Organic farming and the sustainability of agricultural systems. Agric. Syst., 68, 21–40.

Riley, M., Sangster, H., Smith, H., Chiverrell, R. & Boyle, J. (2017). Will farmers work together for conservation? The potential limits of farmers’ cooperation in agri-environment measures. Land use policy.

Romero, A., Chamorro, L. & Sans, F.X. (2008). Weed diversity in crop edges and inner fields of organic and conventional dryland winter cereal crops in NE Spain. Agric. Ecosyst. Environ., 124, 97–104.

Roschewitz, I., Gabriel, D., Tscharntke, T. & Thies, C. (2005). The effects of landscape complexity on arable weed species diversity in organic and conventional farming. J. Appl. Ecol., 42, 873–882.

Rose, F. & O’Reilly, C. (2006). The Wild Flower Key: How to Identify Wild Flowers, Trees and Shrubs in Britain and Ireland. Frederick Warne.

Roulston, T.H. & Cane, J.H. (2000). Pollen nutritional content and digestibility for animals. Plant Syst. Evol., 222, 187–209.

Rundlöf, M., Edlund, M. & Smith, H.G. (2009). Organic farming at local and landscape scales benefits plant diversity. Ecography, 18, 303.

Rundlöf, M., Nilsson, H. & Smith, H.G. (2008). Interacting effects of farming practice and landscape context on bumble bees. Biol. Conserv., 141, 417–426.

Sánchez-Bayo, F. & Wyckhuys, K.A.G. (2019). Worldwide decline of the entomofauna: A review of its drivers. Biol. Conserv., 232, 8–27.

Sotherton, N. & Self, M.J. (2000). Changes in plant and arthropod diversity on lowland farmland: an overview. In: Ecology and conservation of lowland farmland birds (eds. Aebischer, N.J., Evans, A.D., Grice, P.V. & Vickery, J.A.). British Ornithologists Union, pp. 26–35.

Steffan-Dewenter, I., Münzenberg, U., Bürger, C., Thies, C. & Tscharntke, T. (2002). Scale-Dependent Effects of Landscape Context on Three Pollinator Guilds. Ecology, 83, 1421–1432.

Tuck, S.L., Winqvist, C., Mota, F., Ahnström, J., Turnbull, L.A. & Bengtsson, J. (2014). Land-use intensity and the effects of organic farming on biodiversity: a hierarchical meta-analysis. J. Appl. Ecol., 51, 746–755.

Weibull, A.-C., Östman, Ö. & Granqvist, Å. (2003). Species richness in agroecosystems: the effect of landscape, habitat and farm management. Biodiversity & Conservation, 12, 1335–1355.

Williams, N.M., Regetz, J. & Kremen, C. (2012). Landscape-scale resources promote colony growth but not reproductive performance of bumble bees. Ecology, 93, 1049–1058.

Wood, T.J., Holland, J.M. & Goulson, D. (2016). Providing foraging resources for solitary bees on farmland: current schemes for pollinators benefit a limited suite of species. J. Appl. Ecol.

Wood, T.J., Holland, J.M., Hughes, W.O.H. & Goulson, D. (2015). Targeted agri-environment schemes significantly improve the population size of common farmland bumblebee species. Mol. Ecol., 24, 1668–1680.

Zurbuchen, A., Landert, L., Klaiber, J., Müller, A., Hein, S. & Dorn, S. (2010). Maximum foraging ranges in solitary bees: only few individuals have the capability to cover long foraging distances. Biol. Conserv., 143, 669–676.

