## Supplementary materials for "Plugging the hunger gap: Organic farming supports more abundant nutritional resources for bees at critical periods"

### Supporting Information

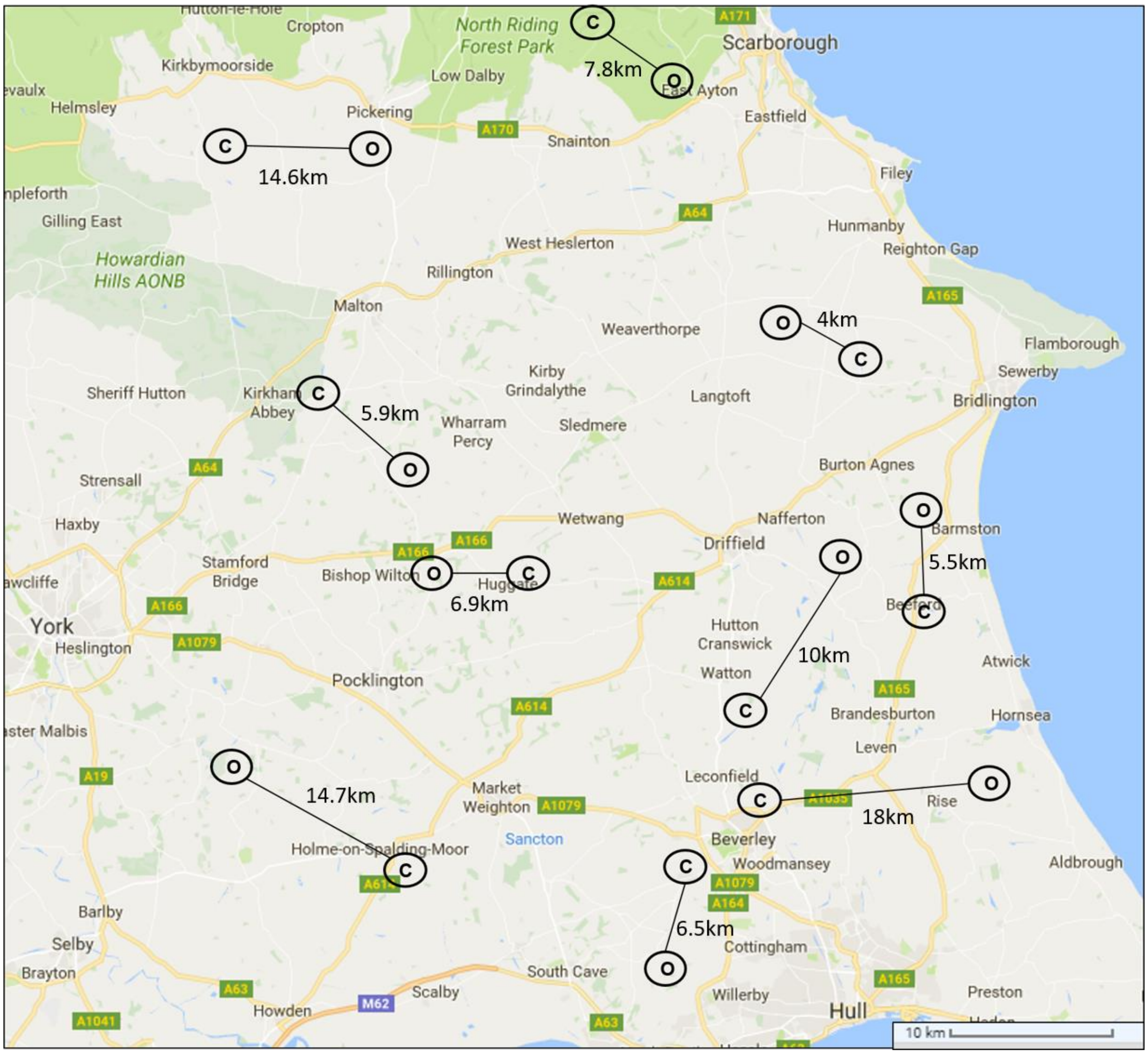


**Fig. S1. Map of farm sites sampled during the study. Letters represent farm type (c = conventional, o = organic) and lines represent site pairings, with approximate distances between farms.**

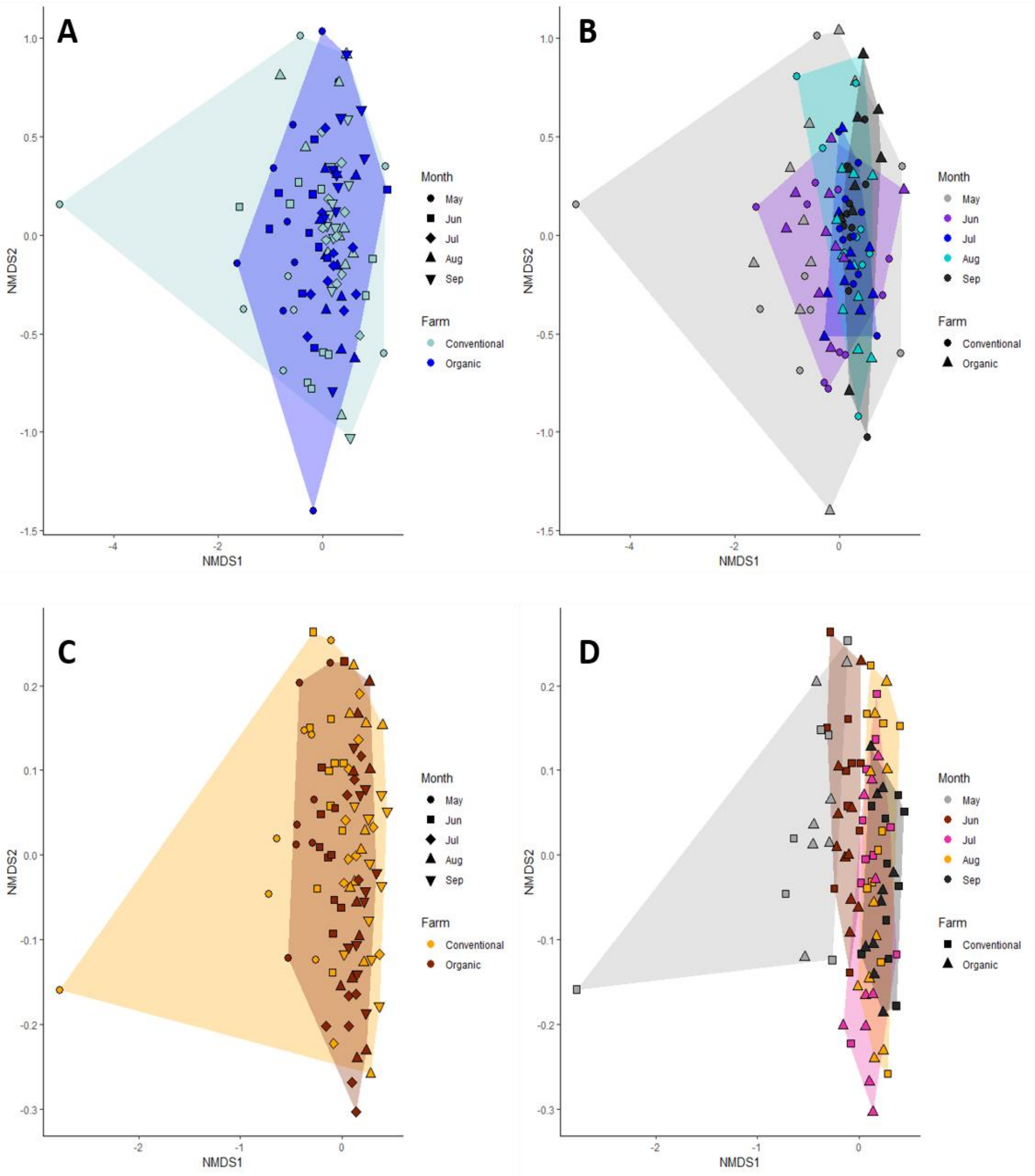


**Fig. S2. NMDS plot for counts of bee (A & B) and plant (C & D) species at each site during each sampling month (converted to percentages), shaded by farm type (A & C) and sampling month (B & D) with the outlying point from the single conventional farm in May included.**

**Table S1. List of individual farm sites with corresponding details of practice, soil types and crops farmed**

| Farm Code | Farm Pair | Farming Practice | Farm Type | Soil Type | Crop |
|---|---|---|---|---|---|
| DEN | P1 | Organic | Mixed | clay - slightly acid | Winter Wheat |
| MLS | P1 | Conventional | Arable | clay - slightly acid | Winter Wheat |
| BES | P2 | Conventional | Arable | clay - slightly acid | Winter Wheat |
| CAR | P2 | Organic | Arable | clay - slightly acid | Spelt Wheat |
| BAR | P3 | Organic | Mixed | clay - slightly acid | Winter Wheat |
| BEE | P3 | Conventional | Arable | clay - slightly acid | Winter Wheat |
| COL | P4 | Organic | Mixed | shallow - chalk/limestone | Winter Barley |
| LIT | P4 | Conventional | Mixed | shallow - chalk/limestone | Winter Barley |
| FRE | P5 | Conventional | Arable | Free-draining loamy | Winter Barley |
| STR | P5 | Organic | Arable | Free-draining loamy | Winter Oats |
| BEC | P6 | Conventional | Arable | Free-draining loamy | Spring Barley |
| BRO | P6 | Organic | Mixed | Free-draining loamy | Spring Oats |
| WAL | P7 | Conventional | Arable | Free-draining loamy | Winter Wheat |
| YOR | P7 | Organic | Arable | Free-draining loamy | Winter Wheat |
| HAS | P8 | Conventional | Arable | Wet sandy loam - acid | Winter Wheat |
| STO | P8 | Organic | Arable | Wet sandy loam - acid | Winter Wheat |
| BXA | P9 | Conventional | Pastoral | Free-draining acid loam | grassland - cattle |
| THO | P9 | Organic | Pastoral | Free-draining acid loam | grassland - cattle |

| | | | | | |
|---|---|---|---|---|---|
| LEA | P10 | Organic | Pastoral | clay - slightly acid | grassland - cattle |
| THR | P10 | Conventional | Pastoral | clay - slightly acid | grassland - cattle |

**Table S2. Names of unique bee taxa according to farming method. Numbers in brackets represent incidences. Note: all these species are considered generalist in their flower usage**

| Conventional only | Organic only |
|---|---|
| *Andrena barbilabris* (1) | *Andrena minutula* (1) |
| *Lassioglossum morio* (2) | *Coelioxys rufescens* (1) |
| *Nomada ruficornis* (1) | *Hylaeus communis* (1) |
| *Sphecodes gibbus* (1) | *Megachile willughbiella* (1) |
| | *Melecta albifrons* (1) |
| | *Nomada leucophthalma* (3) |

**Table S3. Names of unique plant taxa for each farming method. Numbers represent number of floral units**

| Conventional only | Organic only |
|---|---|
| *Ajuga reptans* (3) | *Solanum dulcamara* (126) |
| *Prunus domestica* (105) | *Borago officinalis* (2) |
| *Hypochaeris radicata* (49) | *Arctium minus* (5) |
| *Ranunculus sceleratus* (8) | *Hypochaeris glabra* (1) |
| *Pulicaria dysenterica* (458) | *Cichorium intybus* (60) |
| *Ranunculus arvensis* (4) | *Stellaria media* (1) |
| *Centaurea montana* (22) | *Geranium sp.*(29) |
| *Mentha arvensis* (66) | *Hesperis matronalis* (6) |
| *Petroselinum segetum* (67) | *Vicia villosa* (52) |
| *Agrostemma githago* (10) | *Geranium columbinum* (17) |
| *Rorippa sp.* (3) | *Sonchus asper* (29) |
| *Cardamine pratensis* (15) | *Odontites vernus* (56) |
| *Bellis perennis* (963) | *Matricaria recutita* (208) |
| *Thlaspi arvense* (7) | *Capsella bursa-pastoris* (44) |
| *Veronica chamaedrys* (20) | *Ranunculus parviflorus* (3) |
| *Pentaglottis sempervirens* (11) | *Crepis capillaris* (77) |
| *Crepis sp.*(2) | *Epilobium tetragonum* (30) |
| *Calystegia sepium* (91) | *Stellaria crassiflora (*55) |

*Lamium amplexicaule* (7)

*Geranium robertianum* (135)

*Trifolium dubium* (315)

*Solanum physalifolium* (6)

*Epilobium roseum* (41)

*Primula vulgaris* (10)

*Lychis flos-cuculi* (17)

*Chaerophyllum temulum* (100)

*Leontodon hispidus* (98)

*Geranium pusillum* (20)

*Achillea ptarmica* (3)

*Rubus saxatilis* (1)

*Geum urbanum* (2)

*Mycelis muralis* (75)

*Raphanus raphanistrum* (1383)

*Galium odoratum* (29)
